# Both transient and sustained MPK3/6 activities positively control expression of NLR genes in PTI and ETI

**DOI:** 10.1101/2021.04.01.437866

**Authors:** Julien Lang, Baptiste Genot, Jean Bigeard, Jean Colcombet

## Abstract

*Arabidopsis thaliana* Mitogen Activated Protein Kinases 3 and 6 (MPK3/6) are known to be activated transiently in PAMP-Triggered Immunity (PTI) and durably in Effector-Triggered Immunity (ETI). However the functional differences between these two kinds of activation kinetics and how they allow coordination of the two layers of plant immunity remain poorly understood. Here, by analysing suppressors of the phenotype caused by a constitutively active form of MPK3, we demonstrate that ETI-mediating nucleotide-binding domain leucine-rich repeat receptors (NLRs) and NLR signaling can act downstream of MPK3 activities. Moreover we provide evidence that both sustained and transient MPK3/6 activities positively control the expression of at least two NLR genes, AT3G04220 and AT4G1110. We further show that the ETI regulators NDR1 and EDS1 also contribute to the upregulations of these two NLRs not only in an ETI context but also in a PTI context. Remarkably, while in ETI, MPK3/6 activities are dependent on NDR1 and EDS1, they are not in PTI, suggesting that if the same actors are involved in the two layers of immunity, the way they are interconnected is different. Finally we demonstrate that expression of the NLR AT3G04220 is sufficient to induce expression of defense genes from the SA branch. Overall this study enlarges our knowledge of MPK3/6 functions during immunity and gives a new insight into the intrication of PTI and ETI.

## Introduction

The plant defense responses to pathogens are usually viewed as a two-layered system ^1^. In the first one, cell surface-localized Pattern-Recognition Receptors recognize conserved Pathogen-Associated Molecular Patterns (PAMPs), thereupon eliciting PAMP-Triggered Immunity (PTI) ^2^. In the second layer, pathogen effectors, secreted to counteract plant defense responses and to favor plant susceptibility, are recognized by intracellular Nucleotide-Binding domain Leucine-rich repeat Receptors (NLRs) giving rise to the Effector-Triggered Immunity (ETI) ^3^. If signaling events involved in PTI are relatively well known including Receptor-Like Cytoplasmic Kinases (RLCKs), Mitogen-Activated Protein Kinases (MAPKs), Calcium-Dependent Protein Kinases (CDPKs) and ROS ^4^, the signaling mechanisms of ETI, on the other hand, remain more elusive. The existence of two main classes of NLRs, containing in their N-terminal part either a Coiled-Coil (CC) domain for CC-NLR (CNL) or a Toll and Interleukin-1 Receptor (TIR) domain for TIR-NLR (TNL), suggests that ETI signaling might be processed through two distinct pathways depending on the type of NLR involved. For instance initial reports supported the notion that Non-race specific Disease resistance 1 (NDR1) gene mediates CNL-ETI while Enhanced Disease Susceptibility 1 (EDS1) gene contributes to TNL-ETI ^5^. However further reports undermined this conception, by revealing that CNL-ETI could be independent of NDR1 ^6,7^, that EDS1 could play a role in CNL-ETI ^8,9^, and that CNLs and TNLs could cooperate ^10^.

In addition to this, the strict dichotomy between PTI and ETI that would be consecutive in time and would each represent a specific kind of immunity, has been regularly challenged. Studies showing PTI and ETI not only share numerous signaling components but also lead to similar gene reprogramming progressively built a model in which PTI and ETI are continuously linked, with connections allowing sophisticated and extensive modulations of plant defense responses ^11–14^. In this context it has been recently uncovered that PTI responses enable optimization of ETI responses and that in return ETI responses promote accumulation of PTI actors, demonstrating thereby a mutal potentiation of the two kinds of immunity ^15,16^. Despite these significant advances, the question of the molecular mechanisms underlying the crosstalks between PTI and ETI remains one of the most exciting in the field of plant-microbe interactions ^17^.

MAPKs are essential signaling components that allow plants to integrate various cues coming from biotic and abiotic stresses or developmental programs, into appropriate cell responses ^18^. A canonical MAPK cascade encompasses a MAPK Kinase Kinase (MAP3K), a MAPK Kinase (MAP2K or MKK) and a MAPK (or MPK) which activate in a serial manner by phosphorylation ^19^. Active MAPKs can subsequently phosphorylate specific substrates on specific sites, thereby translating signal inputs into functional outputs ^20^. In the context of immunity, two MAPK cascades have been particularly well characterized. Thus the rapid and transient activation of both MAP3K3/5-MKK4/5-MPK3/6 and MEKK1-MKK1/2-MPK4 cascades, upon PAMP perception, leads to a gene reprogramming that is instrumental in mounting succesful PTI responses ^21–24^. Recently it has been shown that MPK3/6 could also be activated in a sustained manner in response to effector recognition, and several roles, such as buffering of the SA sector of defense, promotion of camalexin production, or inhibition of the photosystem have been associated with this phenomenon ^25–27^. Nonetheless it is not clear so far whether these functions are specific of sustained MPK3/6 activations or the simple extension of processes already controlled by transient MPK3/6 activations. Similarly the question whether sustained MPK3/6 activations constitute a general feature of ETI or are restricted to peculiar effector/NLR recognitions remains debatable ^28^.

Here, starting with an analysis of the regulation and functions of sustained MPK3/6 activations during ETI, we came to the finding that MPK3/6 activities could bridge PTI and ETI by positively controlling the expression of some NLR genes. Combining different genetic backgrounds as well as various pathogen treatments that trigger selective patterns of MPK3/6 activations, we notably provided evidence that the expression inductions of the two NLRs AT3G04220 and AT4G11170 are dependent on both transient and sustained MPK3/6 activities in response to PAMP and pathogen effectors. We also showed that ETI-regulating NDR1 and EDS1 are involved in this process. Altogether our results unveil an original intrication between PTI and ETI components.

## Results

### Sustained MAPK activation is characteristic of RPS2/RPM1-mediated ETI responses, concerns mostly MPK3 and leads to a nuclear accumulation of MPK3

Sustained MPK3/6 activities have been reported in response to pathogen effectors ^25,27,29^. Yet whether these activations represent a general feature of ETI remains controversial ^28,30,31^. To get a better understanding of the question, we decided to compare the pattern of MPK3/6 activities at early (1hpi) and late (8hpi) timepoints, in Col-0 plants and after infiltration with various *Pseudomonas syringae* pv. *tomato* DC3000 strains, expressing different effectors (WT, AvrRpt2 and AvrRps4) or impaired in effector translocation (*hrcc*-). As shown in Figure 1A, all tested strains could induce MPK3/6 activities 1hpi, suggesting that at this early timepoint PAMPs of all the strains or Damage-Associated Molecular Patterns produced by the infiltration can be recognized by the plants and trigger defense signaling. In contrast at later timepoints, MPK3/6 activities were drastically reduced in the samples infiltrated with DC3000 WT, AvrRps4 and *hrcc*-comparatively to the ones infiltrated with AvrRpt2.

**Figure 1:**
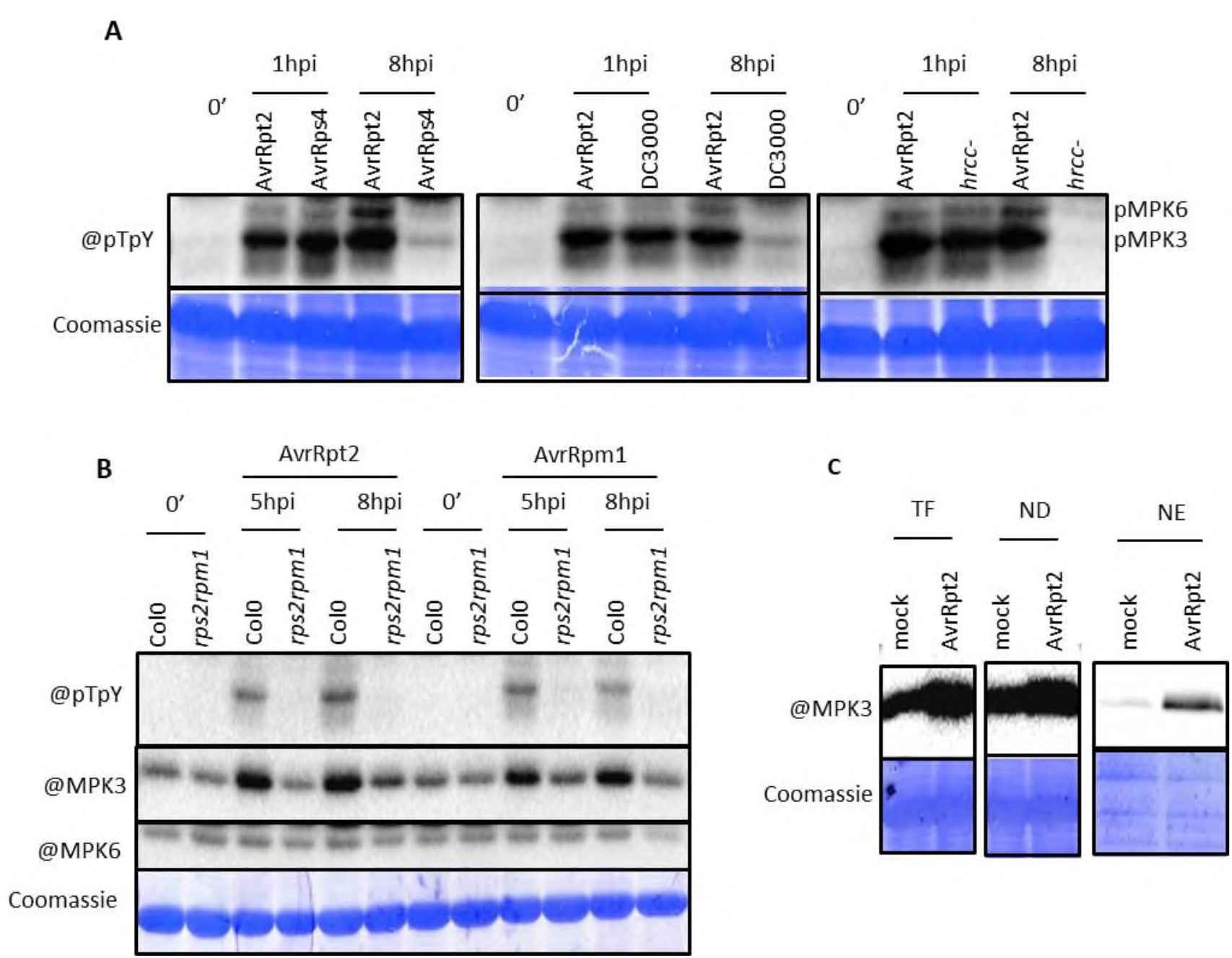
Sustained MAPK activation is characteristic of RPS2/RPM1-mediated ETI responses, concerns mostly MPK3 and leads to a nuclear accumulation of MPK3. MPK3/6 activities and quantities were analyzed through anti-pTpY, anti-MPK3 and anti-MPK6 Western-blots in Col-0 in response to various DC3000 strains (**A**), in Col-0 and *rps2rpm1* backgrounds in response to DC3000 AvrRpt2 and AvrRpm1 (**B**), and in total fraction (TF), nuclei-depleted fraction (ND) and nuclei-enriched fraction (NE) from Col-0 plants in response to mock or DC3000 AvrRpt2 (**C**). Coomassie serves as a loading control. Experiments A and B were repeated at least 2 independent times with similar results.

As AvrRpt2 is known to target RIN4, we compared its effects with the ones of AvrRpm1, another effector targeting RIN4 ^32^, both in WT and the *rps2rpm1* background which is not able to recognize AvrRpt2 and AvrRpm1 anymore. Results indicated that both effectors induce a similar activation of MPK3/6 and that this activation is completely abolished in the *rps2rpm1* background (Fig. 1B). Furthermore in all the experiments, we noticed that sustained MAPK activation concerned mostly MPK3 compared to MPK6. Besides sustained MPK3 activation was correlated with a concomitant increase in the amount of MPK3 proteins whereas amounts of MPK6 remained globally unchanged (Fig 1B).

To determine whether sustained MPK3 activity could affect its subcellular localization, we also quantified MPK3 protein abundance in nuclear and cytoplasmic fractions. The results of the Figures 1C and S1 show that both fractions contain more MPK3 in response to AvrRpt2 infiltration than in response to mock infiltration. However the nuclear fraction is considerably more enriched (more than 10 times) than the cytoplasmic fraction (about 2 times), clearly indicating that in response to AvrRpt2 MPK3 accumulates in the nucleus.

### NDR1 and EDS1 contribute to sustained but not transient MAPK activation

Both NDR1 and EDS1 are involved in the ETI signaling caused by AvrRpt2 recognition ^5,6,8,9^. To determine whether there is a link between these two regulators and sustained MPK3/6 activities, we measured the latters in the *ndr1-1* and *eds1-2* backgrounds. As shown in Figures 2A and 2B, there is a significant decrease in MPK3/6 activities in the two mutants, demonstrating that NDR1 and EDS1 act upstream of the MAPKs and contribute to their activation. By quantification of blot signals from independent experiments, we also concluded that the contribution of NDR1 is higher than that of EDS1 (Fig. 2A and 2B). To consolidate this result, we crossed the *ndr1-1* and *eds1-2* lines with the XVE-AvrRpt2 line that allows direct expression of the AvrRpt2 effector in the plant cell through an estradiol-inducible system ^25^ (Fig. S2), and again we could show that sustained MPK3/6 activities elicited by expression of AvrRpt2 are significantly compromised in absence of functional NDR1 and EDS1, with a higher contribution of NDR1 compared to EDS1 (Fig. 2C). Besides, in all these experiments, we noticed that the accumulation of MPK3 is disrupted in the *eds1-2* background (Fig. 2B and 2C), while in the *ndr1-1* background, the accumulation seems impaired when AvrRpt2 is directly expressed in plants (Fig. 2C), but not when AvrRpt2 is delivered by the *Pseudomonas* strain (Fig. 2A).

**Figure 2:**
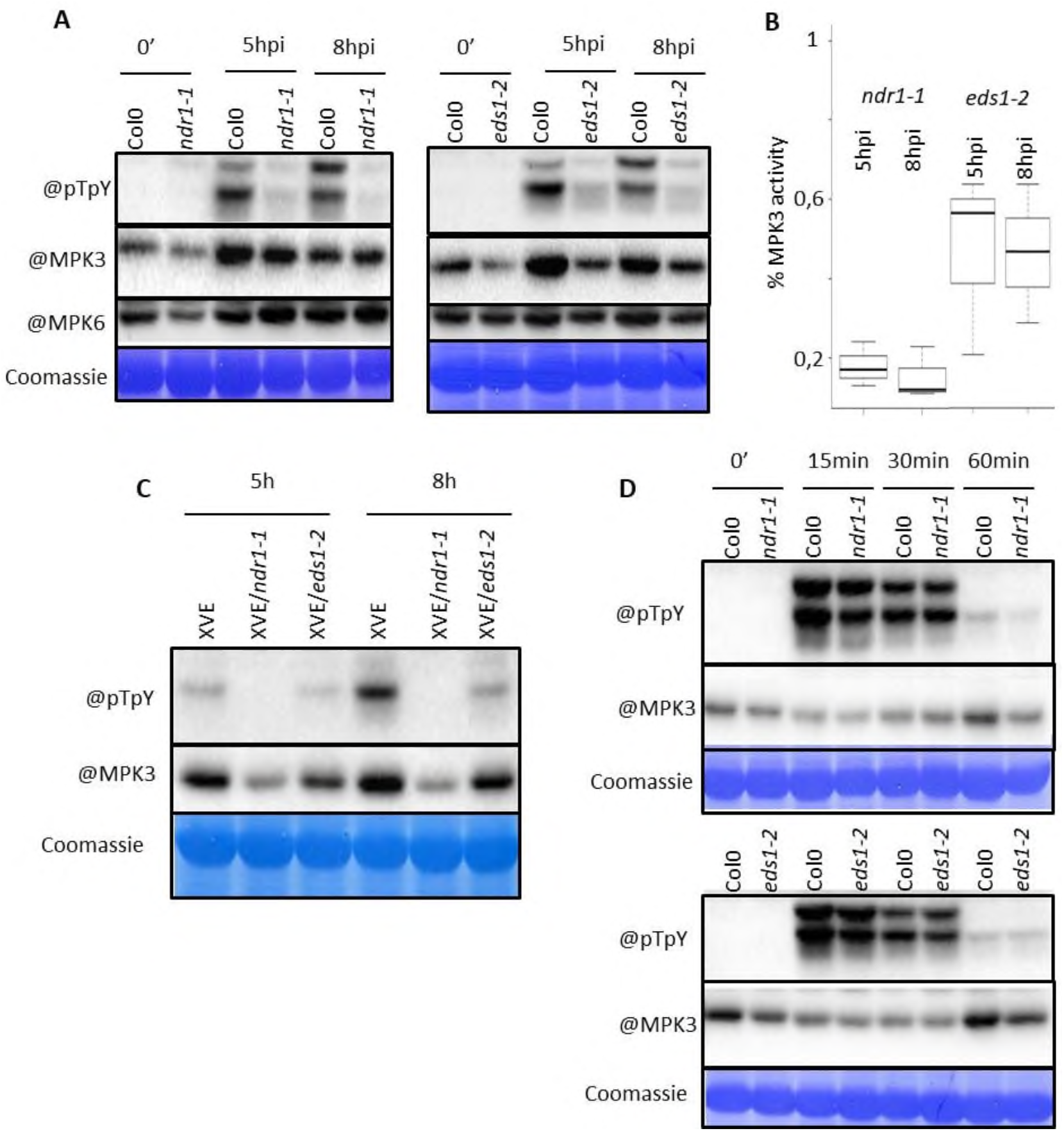
NDR1 and EDS1 contribute to sustained but not transient MAPK activation. (**A**) MPK3/6 activities and quantities were analyzed through anti-pTpY, anti-MPK3 and anti-MPK6 Western-blots in different genetic backgrounds in response to DC3000 AvrRpt2. (**B**) MPK3 activity in *ndr1-1* and *eds1-2* backgrounds comparatively to Col-0. Quantification was performed using ImageJ software from unsaturated Western-blot pictures of 3 independent replicates. (**C**), (**D**) MPK3/6 activities and quantities were analyzed through anti-pTpY, anti-MPK3 and anti-MPK6 Western-blots in different genetic backgrounds in response to estradiol (**C**), and flg22 (**D**). Coomassie serves as a loading control. All experiments were repeated at least 2 independent times with similar results. All experiments were repeated at least 2 independent times with similar results.

Since NDR1 and EDS1 are instrumental in the sustained activation of MPK3/6, we were curious to see whether they also contribute to the transient activation of MPK3/6. To test this we infiltrated Col-0, *ndr1-1* and *eds1-2* leaves with the PAMP flg22 and quantified MPK3/6 activities at early timepoints. However, in this experimental set-up, we could not detect any significant difference between the three genotypes (Fig. 2D), demonstrating that NDR1 and EDS1 are not involved in transient MPK3/6 activations.

### Misregulated MAPK activities result in resistance phenotypes

In an attempt to understand the impact of sustained MPK3/6 activities on the plant defense responses and on the plant resistance to pathogens, we performed pathoassays with *P. syringae* DC3000 AvrRpt2 in different plant backgrounds displaying modifications in the patterns of MPK3/6 activations. The K3CA line is a gain-of-function line that exhibits a higher basal level of MPK3 activity ^33,34^ and also a stronger sustained MPK3 activation upon AvrRpt2 infiltration compared to a K3WT line expressing a WT MPK3 (Fig. S3A). The single mutants *mpk3-1* and *mpk6-4* are defective in the respective MAPKs ^35,36^, yet measurements of their sustained activities upon AvrRpt2 recognition revealed mild effects. As sustained MPK6 activation is weak in response to AvrRpt2, the *mpk6-4* loss of function leaves the strong MPK3 activation unchanged, while in *mpk3-1*, we observed a drastic increase in the levels of MPK6 activations which somehow should compensate for the absence of MPK3 (Fig. S3B and S3C). At last the recently characterized *mkk4mkk5* line ^37^ harbours two weak alleles of the MAP2K genes MKK4 and MKK5 that act directly upstream of MPK3/6 and consistently the *mkk4mkk5* line shows a lower level of MPK3/6 activation both in response to flg22 and AvrRpt2 (Fig. S3D and S3E).

In line with their patterns of MAPK activation, the K3CA line appears more resistant to AvrRpt2 infiltration than WT controls, whereas the *mkk4mkk5* line is more sensitive, and the *mpk3-1* and *mpk6-4* lines behave like Col-0 (Fig. 3). Consistently with their contributions to sustained MPK3/6 activations, we also observed that *eds1-2* is more sensitive than Col-0, in a similar extent as the *mkk4mkk5* line, but less than the *ndr1-1* line (Fig. 3). If these findings tend to suggest that sustained MPK3/6 activities are important for resistance phenotypes, we must remind that the lines we used are not only affected in the patterns of sustained MPK3/6 activations but also in the pattern of transient activations. It is therefore not possible to rule out the possibility that the phenotypes we obtained for the K3CA and *mkk4mkk5* lines are due to defects not in sustained but transient MPK3/6 activations. As a matter of fact, the K3CA and *mkk4mkk5* lines display similar resistance phenotypes in response to DC3000 AvrRps4 infiltration (Fig. S4), even if this effector does not provoke a clear sustained MPK3/6 activation (Fig. 1A).

**Figure 3:**
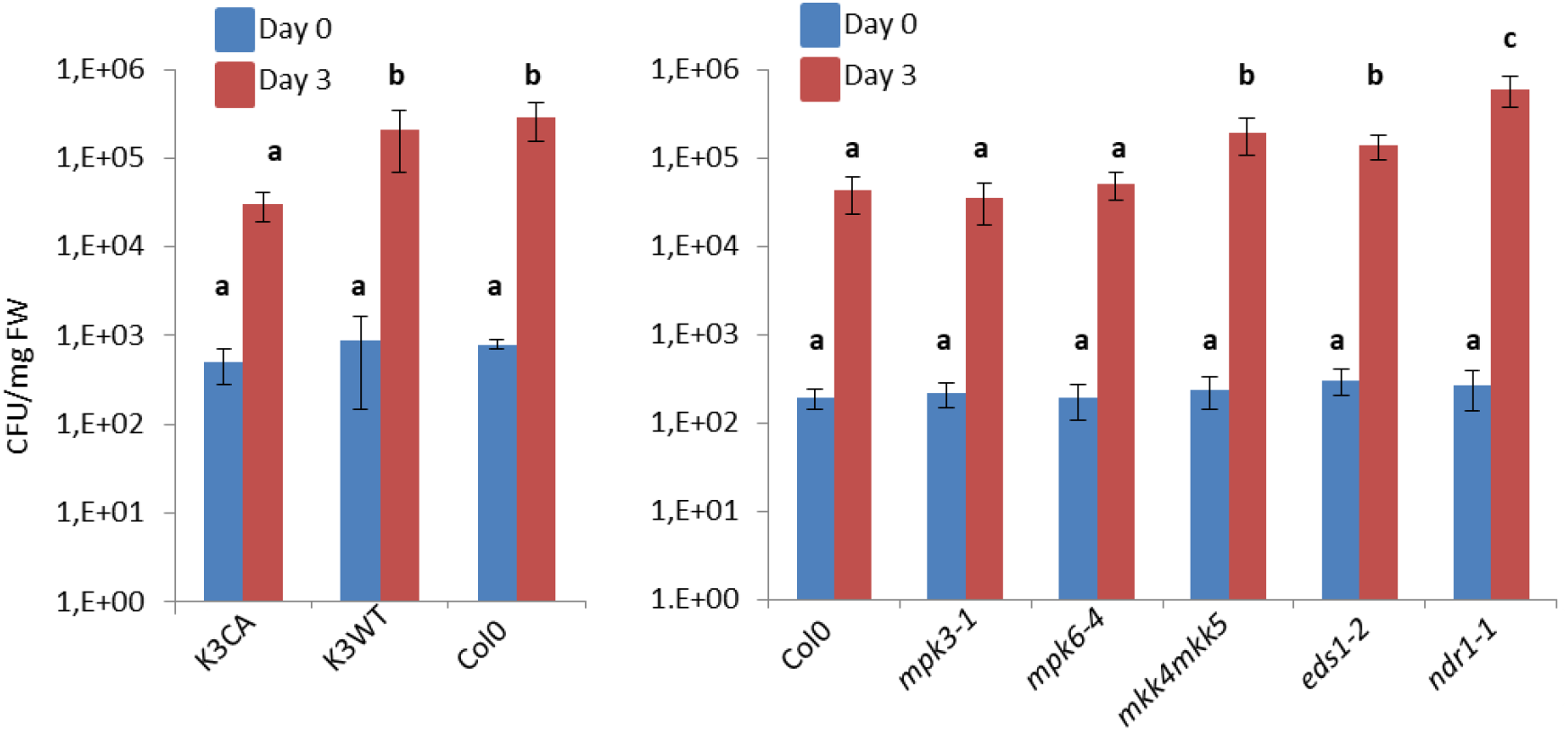
Resistance phenotypes against the DC3000 AvrRpt2 strain. CFU means colony forming unit. Mean values and SDs were calculated from 3 different samples. Letters indicate statistical significance (Mann and Whitney test followed by a Tuckey post-hoc test, p<0,05, n=3). Experiments were repeated 3 times with similar results.

### NLR and NLR signaling act downstream of MPK3 activation

The finding that MPK3 accumulates in the nucleus in response to AvrRpt2 prompted us to look at the genes whose expressions are controlled by MPK3/6 activity. In previous works, we already established that expression of K3CA leads to the upregulation of numerous NLR genes and assumed that these upregulations could be responsible for the auto-immune phenotype of K3CA ^34^. This hypothesis was partly confirmed by the fact that mutation in the CNL SUMM2 partially reverts the K3CA phenotype ^33^. To go further, we crossed K3CA with the *ndr1-1* and *eds1-2* lines as well as with the *snc1-11* line which is impaired in the functions of SNC1, a TNL upregulated in the K3CA transcriptome ^33,34^. As shown in figure 4A, the developmental phenotype of K3CA is partially reverted by the *snc1-11* and *ndr1-1* mutations, and totally by the *eds1-2* mutation. In agreement with this, the accumulations of several SA defense gene transcripts in K3CA are mildly reduced by the *snc1-11* and *ndr1-1* mutations, and drastically by the *eds1-2* mutation (Fig. 4B). Remarkably, when we analyzed the levels of MPK3 activity and quantity in the different lines, we observed that those remain globally the same in K3CA, K3CA/*snc1-11* and K3CA/*ndr1-1*, but considerably decrease in *K3CA/eds1-2* (Fig. 4C). Overall these findings confirm that NLR and NLR signaling can act downstream of MPK3 activity. They also highlight the important role of EDS1 in the regulation of MPK3, especially in allowing high and sustained accumulation of active MPK3.

**Figure 4:**
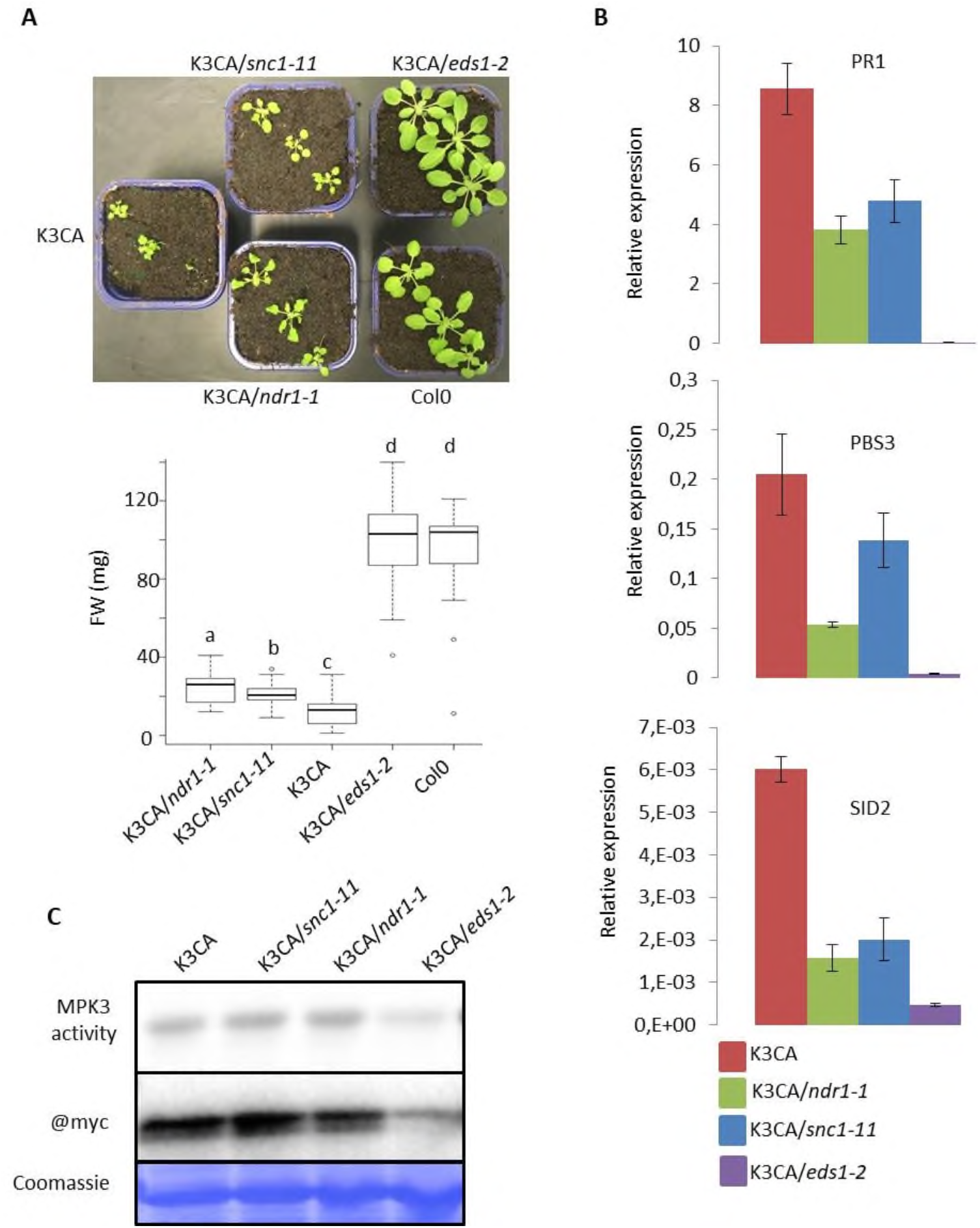
NLR and NLR signaling act downstream of MPK3 activation. (**A**) Developmental reversion of the K3CA phenotype by *ndr1-1*, *snc1-11* and *eds1-2*. Representative picture of 1.5 month old plants (upper part), fresh weight (FW) mass of 1.5 month-old plants (lower part). Letters indicate statistical significance (Mann and Whitney test, followed by a post-hoc Tuckey test, p<0,05, n=19). (**B**) RT-qPCR experiments showing the relative expression levels of SA-related PR1, PBS3 and SID2 genes in different genetic backgrounds in 1.5 month-old plants. Mean values and SDs were calculated from 3 independent replicates. (**C**) K3CA activity and quantity in different genetic backgrounds in 1.5 month-old plants were measured through an anti-myc immunoprecipitation followed by a kinase assay, and an anti-myc Western-blot. Coomassie serves as a loading control. Experiments were repeated at least 2 independent times with similar results.

### The NLRs AT3G04220 and AT4G11170 are upregulated both in ETI and PTI in a manner which is dependent on MPK3/6 activities, EDS1 and NDR1

To get a deeper understanding of the NLR upregulations mediated by MPK3/6 activities, we compared our candidate NLR genes from the K3CA transcriptome ^33,34^ with genes upregulated by constitutive active forms of MKK4/5 ^25,27^, as well as by PAMP treatment ^38^ and by the effector AvrRpt2 ^39^ that cause a transient and sustained activation of MPK3/6 respectively. We ended up with a list of seven NLRs, including five TNLs and two CNLs (Table S1), representing genes likely regulated by MPK3/6 during both PTI and ETI.

Next we performed an expression analysis for two NLRs of our list (AT3G04220 and AT4G11170) in response to various treatments (infiltrations with DC3000 WT, AvrRpt2, AvrRpm1, AvrRps4, *hrcc*-, and mock) at 5hpi and 8hpi corresponding to timepoints where DC3000 AvrRpt2 and AvrRpm1, unlike other strains, induce high sustained MPK3/6 activations. In parallel we also measured the expression levels of the PR1 gene as a readout of the plant SA defense responses. Results revealed that the three genes are strongly upregulated by AvrRpt2 and AvrRpm1, and to a lesser extent by AvrRps4 and *hrcc*-, while the expression levels in response to DC3000 WT and mock remain globally identical (Fig. 5A). From this, we infered that the differences in the AT3G04220, AT4G11170 and PR1 inductions might be mostly due to the differences in sustained MPK3/6 activations.

**Figure 5:**
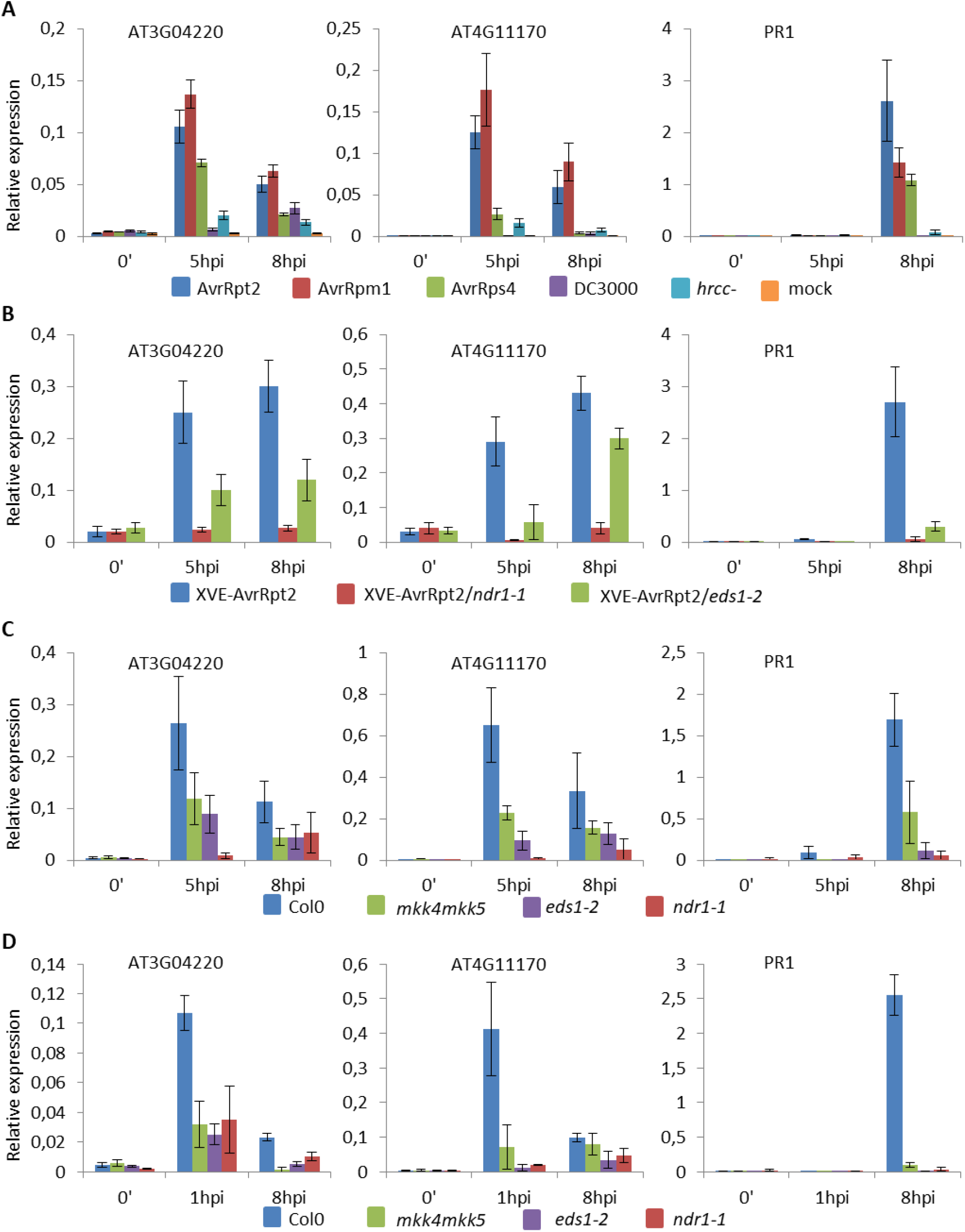
Upregulation of AT3G04220, AT4G11170 and PR1 is dependent on MPK3/6, NDR1 and EDS1. Relative expression levels were measured through RT-qPCR experiments in Col-0 in response to various DC3000 strains (**A**), in different XVE-AvrRpt2 lines in response to estradiol (**B**), in different genetic backgrounds in response to AvrRpt2 infiltration (**C**), and in different genetic backgrounds in response to flg22 (**D**). Mean values and SDs were calculated from 3 independent replicates.

To consolidate our interpretation, we investigated the expression levels of AT3G04220, AT4G11170 and PR1 in the XVE-AvrRpt2 lines. As shown in Fig. 5B, the three genes are strongly upregulated upon estradiol treatment while the inductions are fully compromised in the *ndr1-1* background and partially in the *eds1-2* background. Given the contributions of NDR1 and EDS1 to the activations of MPK3/6 in response to AvrRpt2 (Fig. 2C), these results are consistent with the notion that MPK3/6 activities, NDR1 and EDS1 act in the same signaling pathway to promote expression of AT3G04220 and AT4G11170 during AvrRpt2-triggered ETI.

Furthermore we compared the transcript levels of AT3G04220, AT4G11170 and PR1 in response to AvrRpt2 in the Col-0, *mkk4mkk5*, *ndr1-1* and *eds1-2* backgrounds, and could show that proper induction of the three genes does require functional MKK4/5, NDR1 and EDS1 (Fig. 5C). Similar results were also obtained in response to AvrRpm1 (Fig. S5). Finally we analyzed the expressions of AT3G04220, AT4G11170 and PR1 in the same genetic backgrounds, this time in response to the PAMP flg22. Surprisingly, in this condition, we established that loss of functions of not only MKK4/5, but also of NDR1 and EDS1 impair the upregulation of the NLR genes as well as that of PR1 (Fig. 5D).

### Upregulation of AT3G04220 is sufficient to activate the SA sector of defense

In order to determine what the effects of NLR upregulation can be, we first performed some pathoassays in *at3g04220* and *at4g11170* lines comparatively to Col-0 in response to *P. syringae* DC3000 WT. However we could not detect any significant differences in the load of pathogens between the different genotypes (data not shown), although other studies succeeded in showing that *at4g11170* is more sensitive than WT ^40^.

As an alternative we created two independent transgenic *Arabidopsis* lines expressing the coding sequence of AT3G04220 in the *at3g04220* background, under the control of a dexamethasone-inducible promoter (DEX-At3g04220/*at3g04220* lines). RT-qPCR experiments revealed that the two lines allow a leaky expression of the NLR but still specifically respond to dexamethasone treatment with an induction of more than ten folds (Fig. 6). Then we measured in these two lines and the *at3g04220* mutant, the expression levels of PR1 and PBS3, a gene involved in the synthesis of SA, in response to dexamethasone or mock. With this, we could show that induction of AT3G04220 is correlated with the high induction of the 2 SA marker genes, in a manner which seems dose-dependent (Fig. 6), strongly indicating that control of AT3G04220 expression levels is critical to modulate SA-related defense responses. Interestingly a recent study revealed that ectopic expression of both AT3G04220 and AT4G11170 in tobacco leaves lead to important accumulation of SA and cell death ^41^, which is consistent with our own results.

**Figure 6:**
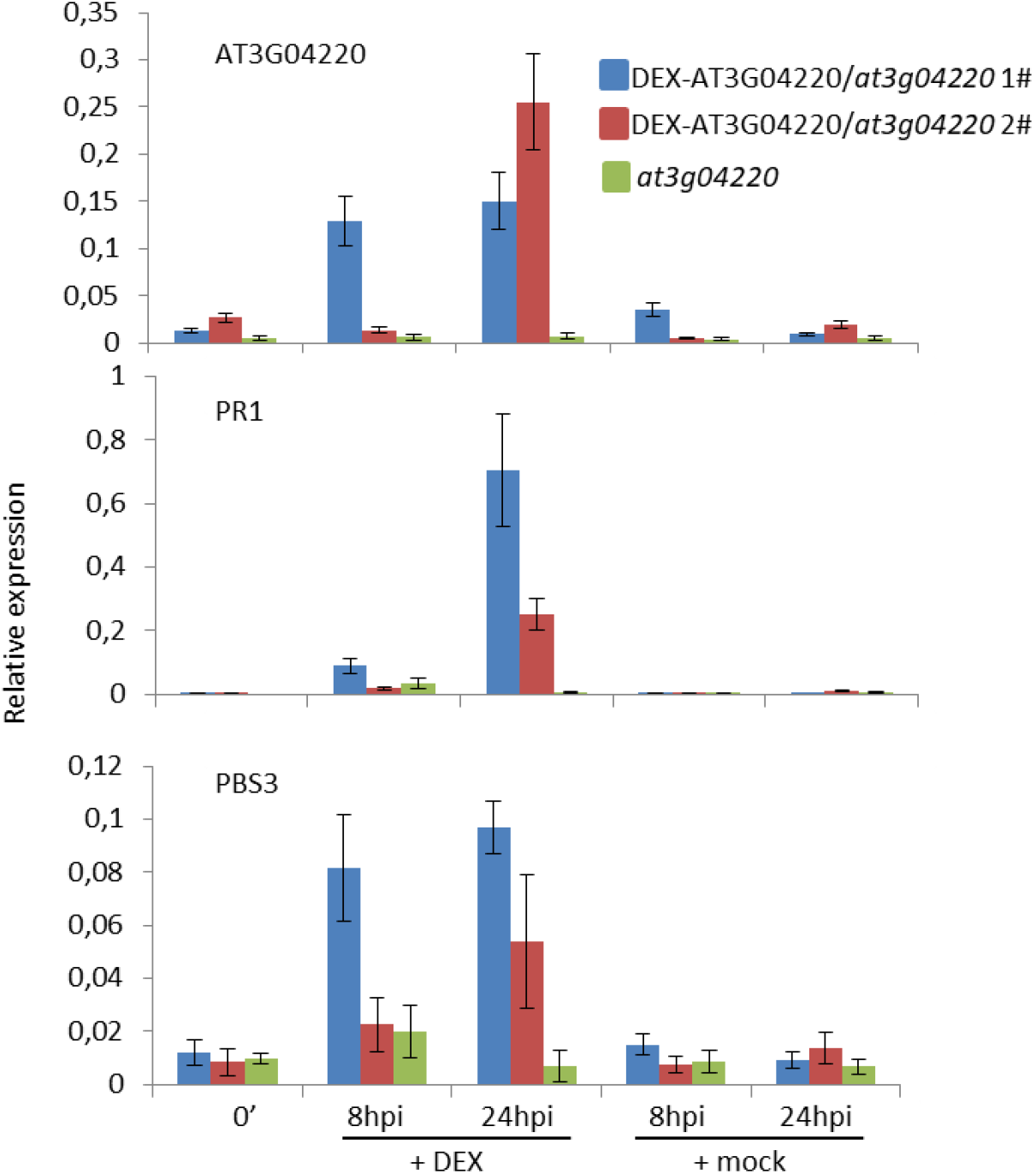
Upregulation of At3g04220 is sufficient to activate SA sector of defense. Relative expression levels were measured through RT-qPCR experiments in 2 independent DEX-AT3G04220/*at3g04220* lines and the at3g04220 line at different timepoints and in response to dexamethasone (+DEX) or mock. Mean values and SDs were calculated from 3 repetitions. The experiment was repeated 2 independent times with similar results.

## Discussion

### Regulation of sustained and transient MPK3/6 activations

One of the first results we obtained by comparing the effects of different *P. syringae* strains expressing different effectors is that sustained activations of MPK3/6 are likely not a general feature of ETI, but rather a characteristic of ETI mediated by the CNLs RPS2 and RPM1 (Fig. 1A,B). This is actually consistent with previous reports ^25,27,30,31^. An explanation could be that sustained MPK3/6 activities are only mediated by CNLs. Another possibility could be that sustained MPK3/6 activities are consecutive to the recognition of effectors acting at the level of the cell membrane, as it is the case for AvrRpt2 and AvrRpm1 which target RIN4. In this model the mechanisms of sustained MPK3/6 activations would be reminiscent of those allowing transient activations ^28^. In the future it would be interesting to analyze the effects on MAPK activation of other effectors, either recognized by CNLs or affecting RIN4 like AvrB ^32^.

We also demonstrated in our study that sustained MPK3/6 activations depend on NDR1 and EDS1 (Fig. 2A, 2B, 2C). Because NDR1 is an integrin-like protein involved in the association between the plasma membrane and the cell wall, as well as a master regulator of the AvrRpt2- and AvrRpm1-triggered ETI ^42^, it is not really surprising to find it upstream of sustained MPK3/6 activations. In AvrRpt2-triggered ETI, EDS1 is known to buffer the SA sector of defense ^8,9^, yet the underpinning molecular mechanisms remain enigmatic. Here our results suggest that EDS1 acts upstream of sustained MPK3/6 to fulfill this function, which is actually in line with the fact that sustained MPK3/6 activities can also buffer the SA sector of defense ^25^.

Three additional points are worth mentioning in regard of the regulation of MPK3/6 activations. First, EDS1 and NDR1 are not required for transient MPK3/6 activations (Fig. 2D), indicating that if both transient and sustained MPK3/6 activations could originate at the cell membrane, there are some decisive discrepancies in the molecular mechanisms of these two phenomena. Second, the *ndr1-1* mutation partially reverts the K3CA phenotype (Fig. 4), strongly suggesting that the NLR signaling component NDR1 acts both upstream and downstream of sustained MPK3/6 activities. Last, the *eds1-2* mutation compromizes the levels of K3CA abundance and activity (Fig. 4C), hinting that EDS1 might be involved in a positive feedback regulation downstream of MPK3 activations allowing the high and sustainable accumulation of the active kinase. This is actually in line with the model where sustained MPK3/6 activations are achieved through a regulatory loop dependent on Systemic-Acquired-Resistance ^29^.

### Functions of sustained and transient MPK3/6 activations

The question whether sustained MPK3/6 activations can give way to new functions comparatively to transient activations is unclear ^28^. Here we provided evidence that NLR upregulation controlled by MPK3/6 is not imputable to a specific pattern of activation, but that both transient and sustained MPK3/6 activations are proficient in it (Fig. 5). What is still missing is a full understanding of the way the transition between transient MPK3/6 activation, elicited by PAMP perception, and sustained activation, elicited by effector recognition, converges towards the NLR upregulations, and thereby impacts the strenght of the defense responses. This is notably a reason why we could not conclude about the resistance phenotypes of lines affected in the same time in the transient and sustained pattern of MPK3/6 activities (Fig.3, S3, S4). Analyzing supplementary timepoints (e.g. 3hpi) might in the future allow us to capture simultaneously the PTI and ETI responses caused by the different *Pseudomonas* strains used in this study and to see how they are integrated at the level of NLR expression. Likewise analyzing later timepoints should reveal if the differences in NLR expression levels are maintained between strains that cause sustained MPK3/6 activation and those that do not, or if ultimately compensatory mechanisms might prevail, resulting in similar responses.

If MPK3/6 activities positively control the upregulations of some NLR genes, the mechanisms underlying these processes are for the moment unknown. Interestingly the inductions of AT3G04220 and AT4G11170 in response to flg22 have been shown to be regulated by promoter DNA methylation and the actions of WRKY transcription factors (TFs) ^38,40^. Since the functions of WRKY TFs are known to be modulated by MPK3/6, either as direct substrates or through the actions of VQ-domain containing proteins ^43^, further investigations in the links between these different actors seem promising.

### NLR upregulation: a crosstalk between PTI and ETI

Upregulations of NLR genes in a PTI context have already been documented in the past ^38^. Nevertheless a comprehensive analysis of the regulation and consequences of this phenomenon is still lacking. Here we demonstrated that MPK3/6, as well as the NLR signaling components EDS1 and NDR1 contribute to the upregulations of two NLR genes, AT3G04220 and AT4G11170, upon both PAMP and effector treatments (Fig. 5). We further showed that upregulation of AT3G04220 is sufficient to activate the SA sector of defense (Fig. 5A, 5B, 6). Altogether these results substantiate a critical role for some NLRs during PTI. They also support the notion that MPK3/6 activities can bridge PTI and ETI by regulating NLR expression levels, thereby modulating or « priming » NLR activation in response to both PAMPs and effectors. Remarkably a concomitant and independent study obtained similar results and came to similar conclusions ^41^. By revealing that TNL accumulation as well as several TNL signaling components, including EDS1, are required to mount an efficient SA-dependent PTI response, the authors of this study pinpointed the same crosstalk between PTI and ETI as we did. However they interpreted the need of TNL signaling components as a downstream event of TNL activation whereas our own data indicate that the NLR signaling components NDR1 and EDS1 might contribute to NLR activation through upregulation.

Overall our findings bring new perspectives to the emerging model of the plant immune system in which defense responses are extensively and dynamically modulated by diverse interactions between PTI and ETI ^13,14^. They also raise important questions that await answers. One of them deals with the identities of the NLRs upregulated and activated during PTI. Are they different from the NLRs involved in recognition of effectors ? For instance Tian et al. claimed that these NLRs are only TNLs ^41^ while in our study NLR genes upregulated by MPK3/6 activities include CNLs as well (Table S1), even if actually we did not analyze them in detail. Another question that seems of the utmost interest is related to the way components of NLR signaling contribute to NLR upregulations during PTI. For instance our results show that EDS1 and NDR1 do not affect transient MPK3/6 activations during PTI (Fig. 2D), suggesting that, contrary to ETI, MPK3/6 activations on one hand, and EDS1 and NDR1 on the other hand act in independent pathways to regulate NLR expression levels. In the same order of idea, Tian et al. showed that TNL inductions are compromised by inhibitor of calcium signaling, suggesting that CDPKs might as well be involved in these processes ^41^. Besides positive feedback loop downstream of NLR activation and promoting NLR expression is also conceivable. The specificities of these different pathways remain to be elucidated.

## Materials and Methods

### Plasmid constructs

The At3g04220 coding sequence was amplified from Arabidopsis cDNA using the primers CACCATGGATTCTTCTTTTTTAC and GCATTTATAAAACTTCAATCTCTTG. Sequencing revealed a 27bp insertion between nucleotide 1935 and 1963 which was not present in the reference sequence from TAIR. This new sequence was then introduced by digestion/ligation between the XhoI and StuI restriction sites in a dexamethasone-inducible expression vector ^44^.

### Plant materials and growth conditions

All plants from this study are in the Columbia background. The *rps2rpm1* ^45^, *eds1-2* ^46^, *ndr1-1* ^47^, *mpk3-1* ^36^, *mpk6-4* ^35^, *mkk4mkk5* ^37^ and K3CA ^33^ backgrounds were described previously. The *snc1-11* (SALK_116460), *at4g11170* (SALK_007034) and *at3g04220* (GABI_290D03) lines were purchased from the NASC and homozygous plants were selected by genotyping using LBb1.3 (SALK), o849 (GABI), TGGTGATTCCGATTTTCTTCCAC and TCTGTTGCTTTAACCTTTGCTCC (*snc1-11*), TTTAGCGGTCAACACGAAAAC and CCAAAATTGAAAATAGAGAACCC (*at4g11170*), and GTCGTCTTTATCTCTCACGCG and GAAGGGCCTCTTCATAGTTGG (*at3g04220*) primers. The DEX-At3g04220 line was obtained by floral dip, and transformed plants were selected on hygromycin. The K3CA/*ndr1-1*, K3CA/*eds1-2* and K3CA/*snc1-11* lines were obtained by crosses and homozygous plants were selected by genotyping.

All plants were grown in growth rooms at 20°C in short day conditions (8h light / 16h dark) at 60% hygrometry and under a light intensity of approximatively 150μmol m^−2^ s^−1^.

### Plant treatments and bacterial infections

All treatments (chemical and bacterial infections) were performed on 1.5 month-old plants by syringe infiltration. The PAMP flg22 and the steroids estradiol and dexamethasone were used at 1μM, 10μM and 5μM respectively. The *P. syringae* pv. *tomato* DC3000 strains WT, AvrRpt2, AvrRpm1, AvrRps4 ^5^ and *hrcc*-^48^ were described previously. The bacteria were grown on solid NYGA medium (0.5% bactopeptone, 0.3% yeast extract, 2% glycerol, 1.5% agar) and liquid LB medium supplemented with the appropriate antibiotics (50μg/ml rifampycin for WT and *hrcc*-, and 50μg/ml rifampycin + 25μg/ml kanamycin for AvrRpt2, AvrRpm1 and AvrRps4).

For protein and RNA analysis, fresh cultures of bacteria were washed and resuspended in 10mM MgCl_2_ at a final OD_600_=0.015 and then infiltrated. Samples were typically constituted from punches of leaves coming from at least 2 different plants.

For pathoassays, fresh cultures of bacteria were washed and resuspended in 10mM MgCl_2_ at a final OD_600_=0.005 and then infiltrated. Samples were constituted from punches of leaves coming from at least 2 different plants. Bacteria load was quantified by counting the colony forming units.

### Protein Methods

For Western blots, proteins were extracted in a nondenaturant buffer (50mM Tris-HCl pH 7.5, 150mM NaCl, 0.1% NP40, 5mM EGTA, 0.1mM DTT) or in nondenaturant Laccus buffer (15mM EGTA, 15mM MgCl_2_, 75mM NaCl, 1% Tween, 25mM Tris-HCl pH 7.5, 1mM DTT) in presence of inhibitors of proteases and phosphatases, and then quantified by Bradford assay. About 10μg of total proteins were loaded on SDS-PAGE gels. The antibodies used were anti-pTpY (Cell Signaling 4370L), anti-MPK3 (Sigma M8318), anti-MPK6 (Sigma A7104), anti H3 (Abcam Ab1791), anti-PEPC (ThermoFischer 4100-4163) at 1/10 000 dilution.

For immunoprecipitation, proteins were extracted in a Laccus nondenaturant buffer in presence of inhibitors of proteases and phosphatases, and then quantified by Bradford assay. About 100μg of total proteins were mixed with 20μl of sepharose beads (GE Healthcare) and 0.5μl of anti-myc (Sigma C3956) antibody and incubated for 2h with gentle shaking. Then the immunoprecipitates were washed 2 times in SUC1 buffer (50mM Tris-HCl pH 7.4, 250mM NaCl, 5mM EGTA, 5mM EDTA, 0.1% Tween) and 2 times in Kinase buffer (20mM Hepes pH 7.5, 15mM MgCl_2_, 5mM EGTA, 1mM DTT).

For kinase assays, immunoprecipitates were resuspended in 15μl of kinase buffer containing 0.1mM ATP, 1mg.ml^−1^ MBP and 2μCi ATP [γ-33P]. After 30min of reaction at room temperature samples were loaded on SDS-PAGE gel. Then the gels were dried and revealed using a STORM scanner.

For nucleocytoplasmic fractioning, proteins were extracted in Honda buffer (2.5% Ficoll type 400, 5% Dextran MW 35-45k, 0.4M sucrose, 25mM Tris-HCl pH 7.5, 10mM MgCl_2_, 5mM DTT) in presence of inhibitors of proteases and phosphatases. After 15min of incubation on ice in presence of 0.5% Triton X-100, an aliquot corresponding to the total fraction was collected. After centrifugation at 1500g for 5min at 4°C, an aliquot of the supernatant corresponding to nuclei-depleted fraction was collected. After washing with Honda buffer + 0.1% Triton X-100, the pellet was resuspended in Honda buffer and an aliquot corresponding to nuclei-enriched fraction was collected.

### RNA Methods

RNA was extracted using Nucleospin^™^ RNA Plus Kit (Macherey Nagel) according to the manufacturer’s instructions and quantified with a Nanodrop spectrophotometer. Typically 1μg of total RNA was used to perform RT, using SuperScript^™^ II Reverse Transcriptase (Invitrogen) and following manufacturer’s instructions. 10ng of cDNA was used for qPCR on a LightCycler^®^ 480 System (96 wells), using LightCycler^®^ 480 SYBR Green I Master (Roche), and following the manufacturer’s standard instructions. ACT2 (AT3G18780) was used as an internal reference to calculate relative expression. Occasionally SAND (AT2G28390) was used as an internal reference to verify that the results were not biased by the choice of ACT2. The primers used for qPCR are listed in Table S2.

## Author Contributions

JL, through insightful discussions with JC, designed the experimental setups and proceeded the data. JL, BG and JB performed the experiments. JL wrote the manuscript with contributions of JB and JC.

## Aknowledgements

The authors would like to thank Drs. Kenishi Tsuda and Wei Zhang for sharing plant and bacterial materials. This study was supported by the French INRAE.

## Supplementary Tables

**Table S1:** List of NLR genes upregulated by K3CA, MKK4/5, flg22, AvrRpt2 and AvrRpm1.

| Gene | Description |
| --- | --- |
| AT1G12290 | CNL localized at the plasma membrane |
| AT4G11170 | TNL (RMG1) |
| AT3G04220 | TNL |
| AT5G41750 | TNL |
| AT1G66090 | TNL |
| AT1G15890 | CNL |
| AT1G57630 | TIR-domain containing protein |

NLR genes commonly upregulated by K3CA (Genot et al., 2017), constitutively active forms of MKK4/5 (Tsuda et al., 2013, Su et al., 2018), flg22 treatment (Yu et al., 2013), and AvrRpt2 and AvrRpm1 infiltrations (Mine et al., 2017)

**Table S2:**
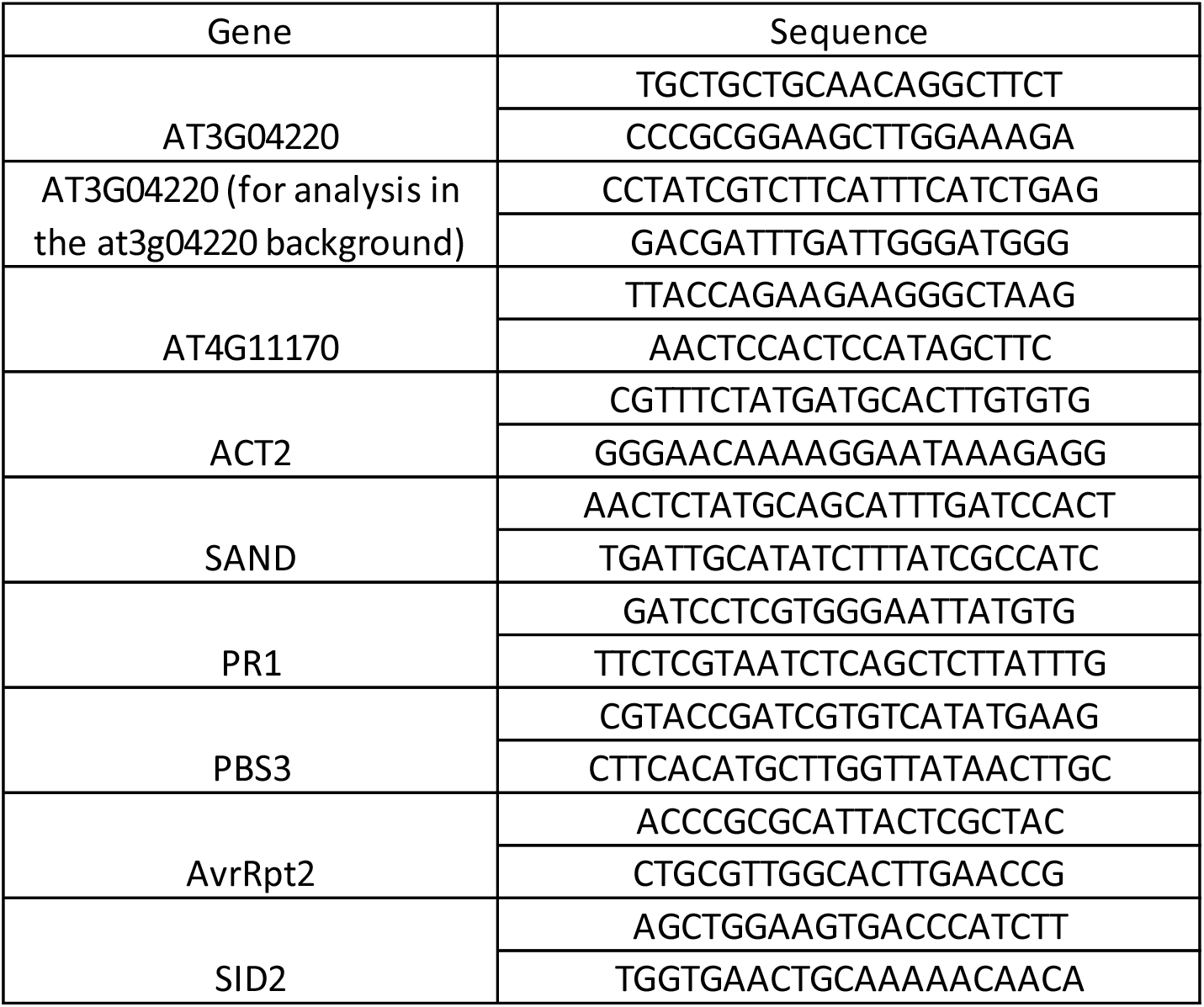
Sequences of primers used for qPCR experiments.

## Supplementary Figure Legends

**Figure S1:**
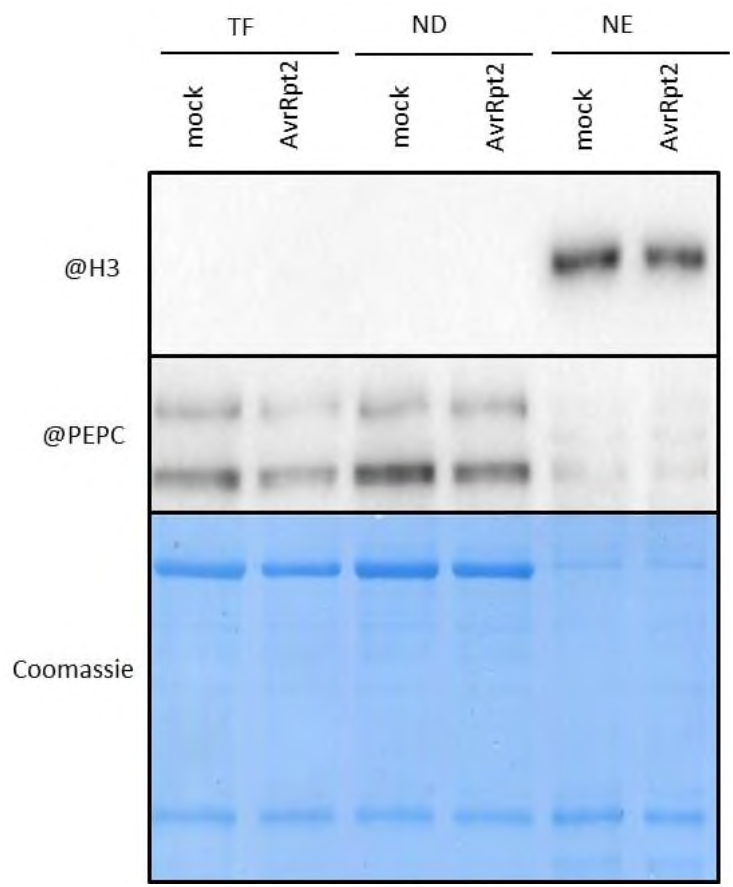
Nucleocytoplasmic fractioning. Anti-Histone 3 (H3) and anti-Phosphoenolpyruvate carboxylase (PEPC) Western-blots as quality markers of the total fraction (TF), nuclei-depleted fraction (ND) and nuclei-enriched (NE) fraction from Col-0 plants infiltrated with mock or DC3000 AvrRpt2. Coomassie serves as a loading control.

**Figure S2:**
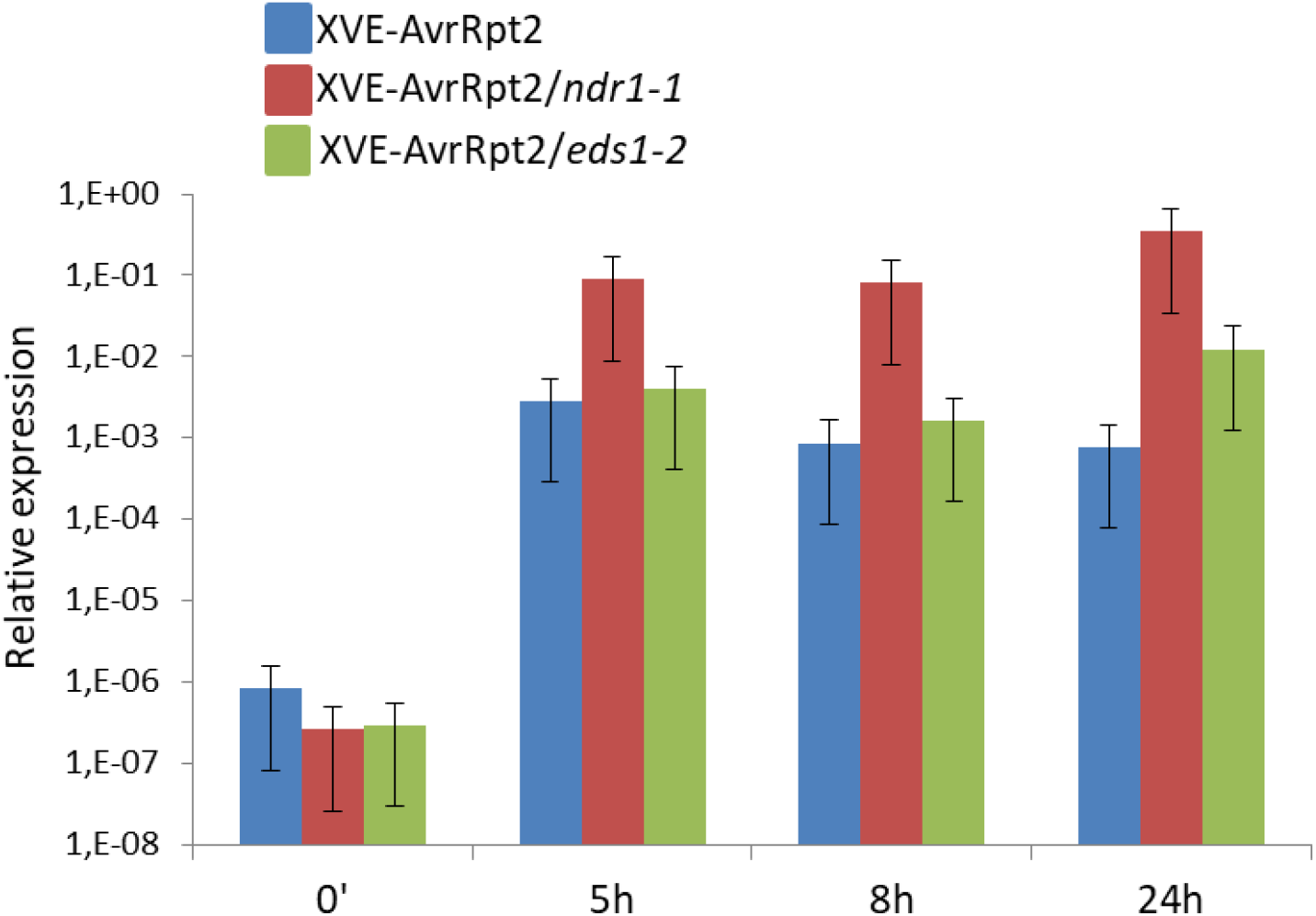
AvrRpt2 induction in the XVE-AvrRpt2 lines. RT-qPCR experiments showing the relative expression levels of AvrRpt2 in the XVE-AvrRpt2 lines at different times after estradiol treatment. Mean values and SDs were calculated from 3 independent replicates.

**Figure S3:**
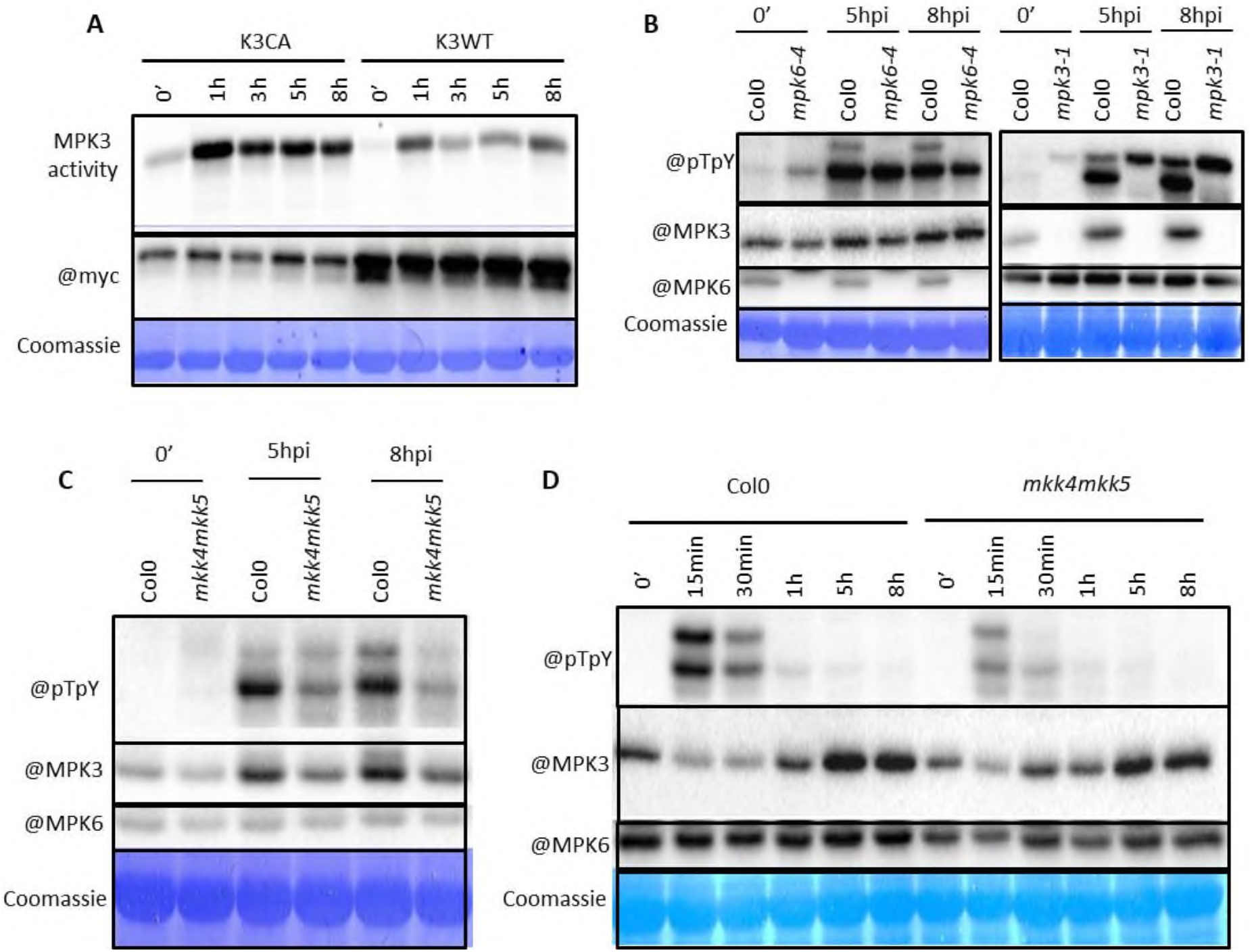
Misregulation of MPK3/6 activities in *K3CA*, *mpk3-1*, *mpk6-4* and *mkk4mkk5* lines. (**A**) K3CA activity and quantity at different times after AvrRpt2 infiltration were analyzed through an anti-myc immunoprecipitation followed by a kinase assay, and an anti-myc Western-blot. (**B**, **C**, **D**) MPK3/6 activities and quantities were measured by anti-pTpY, anti-MPK3 and anti-MPK6 Western-blots at different times after AvrRpt2 infiltration (**B**, **C**) or flg22 treatment (**D**). Coomassie serves as a loading control. All experiments were repeated at least 2 times with similar results.

**Figure S4:**
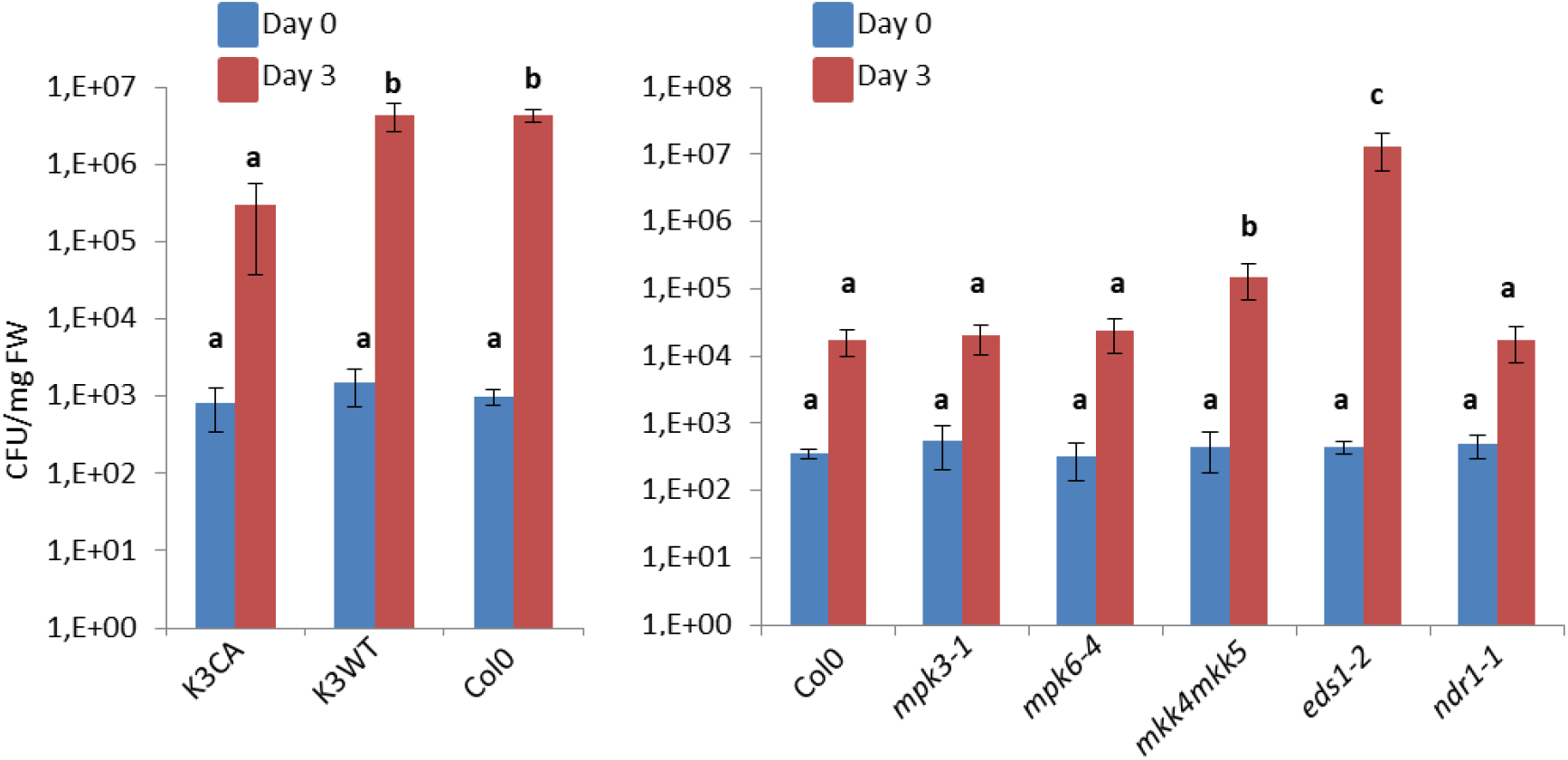
Resistance phenotypes against the DC3000 AvrRps4 strain. CFU means colony forming unit. Mean values and SDs were calculated from 3 different samples. Letters indicate statistical significance (Mann and Whitney test followed by a Tuckey post-hoc test, p<0,05, n=3). Experiments were repeated 3 times with similar results.

**Figure S5:**
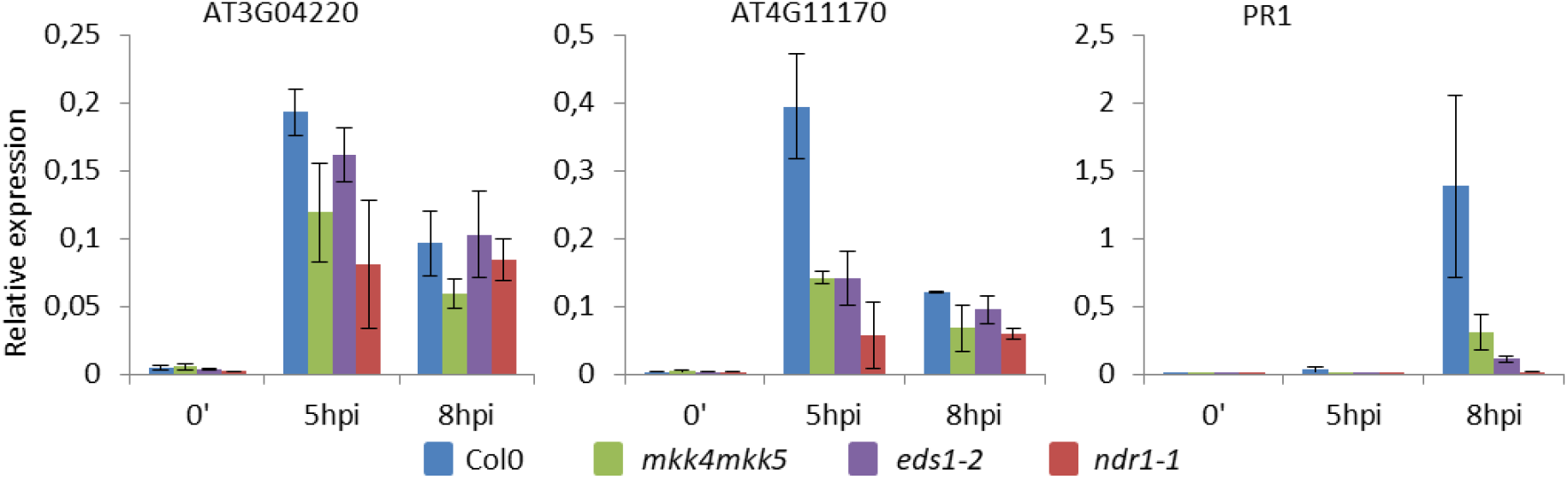
Upregulation of AT3G04220, AT4G11170 and PR1 in response to DC3000 AvrRpm1 infiltration in different genetic backgrounds. Relative expression levels were measured through RT-qPCR experiments. Mean values and SDs were calculated from 3 independent replicates.

